# DeepPlnc: Bi-modal Deep Learning for Highly Accurate Plant lncRNA Discovery

**DOI:** 10.1101/2021.12.10.472074

**Authors:** Ritu, Sagar Gupta, Nitesh Kumar Sharma, Ravi Shankar

**Affiliations:** Studio of Computational Biology & Bioinformatics, The Himalayan Centre for High-throughput Computational Biology (HiCHiCoB, A BIC supported by DBT, Govt. of India), CSIR-Institute of Himalayan Bioresource Technology (CSIR-IHBT), Palampur (HP), 176061, India; Academy of Scientific and Innovative Research (AcSIR), Ghaziabad, Uttar Pradesh, 201002, India

**Keywords:** Plant lncRNAs, Deep learning, Bi-modal CNN, Transcriptomic analysis, Next generation sequencing, non-coding RNAs

## Abstract

We present here a bi-modal CNN based deep-learning system, DeepPlnc, to identify plant lncRNAs with high accuracy while using sequence and structural properties. Unlike most of the existing software, it works accurately even in conditions with ambiguity of boundaries and incomplete sequences. It scored consistently high for performance metrics while breaching accuracy of >98% when tested across a large number of validated instances. During benchmarking it consistently outperformed all the compared tools and maintained a highly significant lead in the range of 4.6%-10.3% from the second best performing tool (p-value << 0.01). DeepPlnc was used to annotate a de novo assembled transcriptome of a himalayan species where again it suggested its much better suitability for genome and transcriptome annotation purposes than the existing tools. DeepPlnc has been made freely available as a web-server and stand-alone program at https://scbb.ihbt.res.in/DeepPlnc/.

## 1. Introduction

~80% of the genome is commonly transcribed into RNA, of which the majority belongs to noncoding RNAs (ncRNAs) alone [1]. More than 80% of these ncRNAs belong to long non-coding RNAs (lncRNAs) [2]. Though not a clear definition or characterization process exists for them, the ncRNAs having length greater than 200 bases are considered as lncRNAs with short open reading frames (<100 amino acids) which are not translated into functional proteins [3]. LncRNAs are supposed to act as a key modulator for various biological processes [4]. Their involvement is reported in controlling transcription process through enhancers and providing regulatory binding sites is well reported [5, 6]. They are also reported to work as sponges to miRNAs and suppress miRNA function by causing deflection in their supposed targeting [7]. The lncRNAs are also found in nucleus and act as major components of nuclear speckles while being a partner in epigenetic and transcriptional controls [8]. In cytoplasm, lncRNAs interact with several RNA binding proteins (RBPs) and control their regulatory dynamics [9].

In plants, the first reporting of lncRNA was done in Soybean for ENOD40 [10]. To this date, the number of experimentally identified lncRNAs in plants is much lesser than those reported for human and mouse, despite the fact that plant genomes are much more complex than animals’ with several genomes much larger than the human genome. Unlike animal system based resources, there is a dearth of such resources for plants. The majority of the entries belong to the predicted ones through some rules based approached and lag behind enormously when compared to animals based resources and information. Seeing such huge dependence of lncRNAs annotations in plants upon software alone itself makes a strong case for research on such tools. And when one looks for computational tools available to identify plant lncRNAs, more so for annotation and identification purpose, there is an enormous scope for improvement.

Compared to animals, plant lncRNAs differ even in their transcriptional modes and display much more complexity and diversity with marked poor degree of sequence conservation. This is one big reason for the evident scarcity of reliable tools and resources for plant lncRNA identification which mostly work with animal system identified properties. Some tools based on regular machine learning approaches like CPC [11] and CPC2 [12] have been developed where the inputs from human datasets were used to train the models. Yet, there are some recent tools like PLncPRO [13], RNAplonc [14], PreLnc [15], and CNIT [16] which have trained models on plant datasets exclusively. Coding Potential Calculator (CPC) and its upgraded version, CPC2, employ Support Vector Machine (SVM) models which depend upon four sequence intrinsic features (Fickett TESTCODE score, ORF length, ORF integrity, isoelectric point) to evaluate the protein-coding potential of transcripts, using ORF prediction quality from a BLASTX search based homology-based approach.

PLncPRO is an another tool based on machine learning approach and applies random forest (RF) algorithm while using 71 features, including BLASTX outputs and frame entropy and frequencies of each of the 64 trimers [13]. Another tool, RNAplonc, uses 16 features including sequence and structural free energy and applied REPTree machine learning algorithm [14]. A more recently published tool, PreLnc, uses incremental feature selection method across the traditionally identified features for lncRNAs. PreLnc is applying Pearson correlation co-efficient to reduce the total features and applies five different classifiers’ comparisons (logistic regression (LR), SVM, decision tree (DT), RF, and K-nearest neighbor methods (KNN)) to build the models [15]. In addition to these tools, CNIT, an upgraded version of CNCI [17] tool, exhibits high accuracy using XGBoost machine learning algorithm. This tool considers both animal and plant datasets for model building. CNIT uses sequence based features such as maximum most-likely CDS score, their lengths, their standard deviation scores, and frequency of 64 codons [16].

Apart from the above mentioned machine learning approaches, which depend a lot upon successful manual identification and features extraction by the authors, very recently efforts have been made to involve deep learning methods to identify lncRNAs. In doing so, such deep learning networks extract important and even hidden transient features which would be otherwise hard to get through manual feature extraction approaches. Deep learning achieves this through intensifying the representation power of complex features by multifold [18]. However, very limited development has happened in this direction for plant lncRNA identification. A tool, PlncRNA-HDeep, has been reported very recently which identifies plant lncRNAs applying two alternative deep learning models, a Long Short Term Memory (LSTM) and a Convolution Neural Net (CNN) framework [19] utilizing predicted dataset only for single species. Although, this tools does not provide any pre-bulit model for classification.

Despite all this progress, a fact remains that there is an enormous scope for improvement for plant specific lncRNA identification. First of all, the existing machine learning approaches now need to move towards deep learning approaches where it would become possible to catch those features which are otherwise difficult to be caught through manual process of feature selection and identification. Traditional features like those based on k-mer and motifs based properties, length, ORF properties, and Kozak sequences may easily fail when sequence boundaries are not clear or shorter incomplete transcripts are reported after assembling. Most of the existing tools report satisfactory scores when tested against some standard benchmarking dataset having full length sequences. Some of these datasets themselves need to be looked into as many instances of them are predicted instances which may lower the quality of training and negatively impact the learning process. As soon as these tools face the real situation where sequence boundaries are not known, be it genomic annotation or annotation of *de novo* assembled transcriptome. Almost all of them fail, raising a big question mark on their actually applicability.

In the present study, we have developed a bi-modal CNN based deep learning model for lncRNAs identification which parallelly worked upon the nucleotides sequences and structural sequences based information. CNNs have proved themselves tremendously successful in detecting spatial arrangements of patterns which are hard to get recognized otherwise. We expected CNN based deep-learning system to successfully detect such spatial arrangements of patterns in nucleotide sequences as well as structural sequences if they were transformed into pixeled form like images. Such approach is more important in real case applications like annotation of genomes and *de novo* assembled transcriptomes where boundaries are not clear and identification needs to be capable enough to work with incomplete sequences. Here, a base frame of 400 bases was used in an overlapping sliding window manner while keeping a cut-off of minimum 200 bases with padding. The entire system was developed by using experimentally validated instances and the system was tested across four different datasets. The developed tool, DeepPlnc, consistently outperformed the compared recently published tools during a highly comprehensive benchmarking process. The study concludes with a practical application demonstration of the software on the *de novo* assembled transcriptome data of a Himalayan medicinal plant where it reported its lncRNAs for the first time.

## 2. Materials and Methods

### 2.1. The datasets

In this study, we considered experimentally validated plant lncRNAs as positive instances and fully annotated non-lncRNA (mRNA) as the negative instances. The lncRNA dataset was collected from Ensembl Plants (v51) with known status, PncStress, and PLncDB V2.0 repositories which provided lncRNA data from 51 plant species. The negative examples, i.e. non-lncRNA (mRNA) were downloaded from Ensembl Plants (v51) for corresponding plant species considered for lncRNAs. On the basis of length, positive and negative datasets were filtered. After filtration, the data was split into train and test sets in the ratio of 50:50. The train datasets were used to build the model and the remaining 50% (test set) was kept untouched during the building of model and were used for performance testing and comparison of the built models. For the performance stability and consistency evaluation, 10-fold independent random trials were performed. Every time, the dataset was split randomly into 50:50 ratio with the first part used to rebuild the model from the scratch and the second part used to test it. In this entire study it was ensured that no instances overlapped between test and train sets in order to ensure unbiased and memory free assessment and model development.

To further evaluate the performance of the DeepPlnc, another dataset was constructed and named as Dataset “B”. The Dataset “B” was derived from 16 plant species. This dataset was based on the known and predicted plant lncRNAs. The predicted lncRNAs were found in the similar manner as was done in previously published studies [12,17]. The source for this dataset was from CNIT which had collected lncRNAs with status as “known” from Ensembl plants whereas the predicted lncRNAs were downloaded from PNRD database which is a comprehensive integrated web resource for ncRNAs (http://structuralbiology.cau.edu.cn/PNRD/). The negative instances (mRNAs) for this dataset were downloaded from Ensembl plants (v51). The sequences were randomly selected for the corresponding 16 species considered for the lncRNA data. All the datasets used in the present study have been provided at the DeepPlnc webserver and in Supplementary Table S1.

Both these datasets were further developed into their corresponding chunked datasets whose details are given below elsewhere. The full illustration of the processing of both the datasets is given in Figure 1.

**Figure 1:**
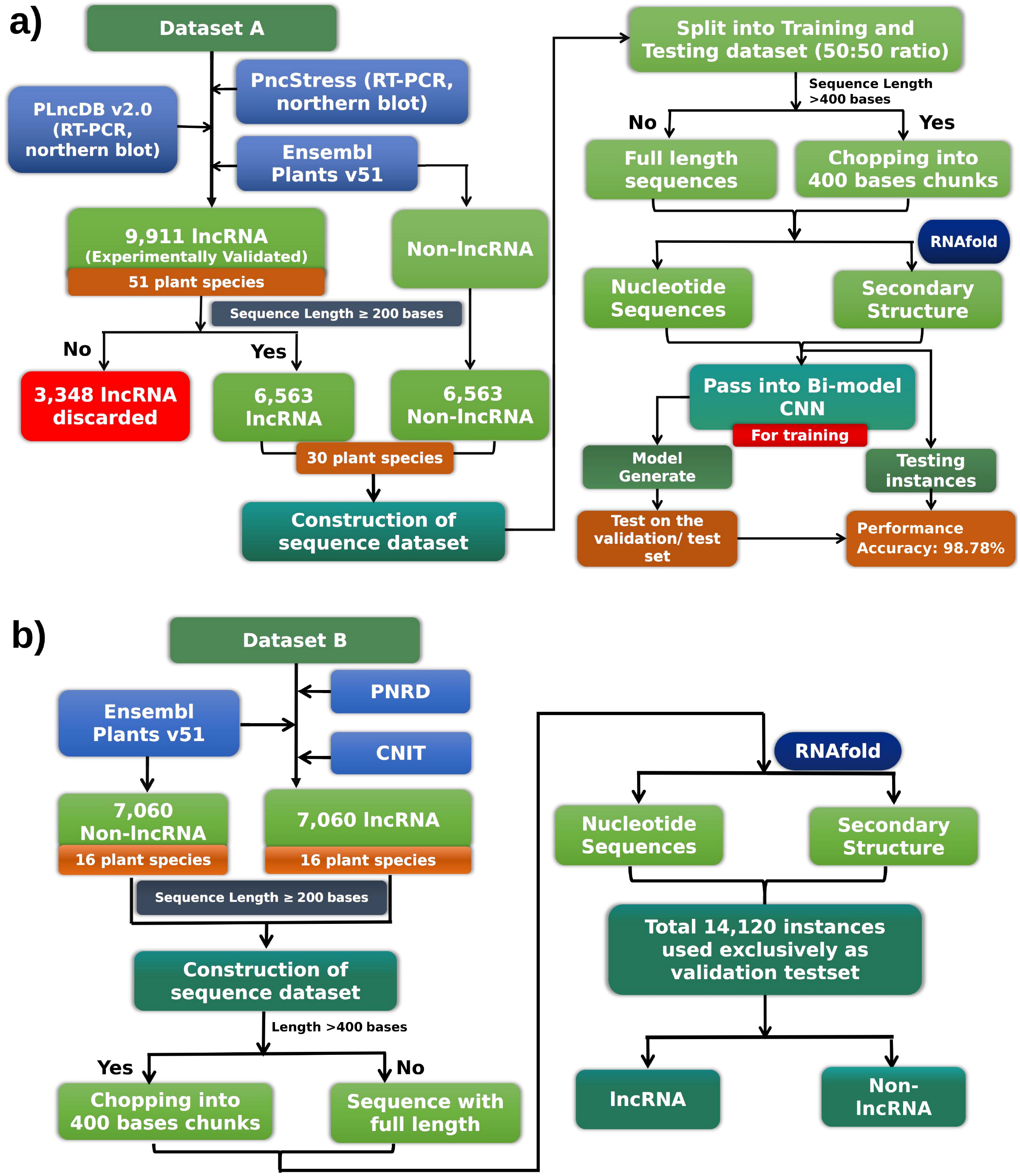
Flowchart representation of dataset processing and formation. **a)** The protocol followed for Dataset A creation, **b)** Protocol followed for Dataset B creation.

### 2.2. One hot encoding and deep-learning implementation

In CNN, the input instances are required to be presented as a tensor of numerical values [28]. Here, we encoded each base in the sequence with “one-hot” encoding (“A”: [1, 0, 0, 0], “T”: [0, 1, 0, 0], “G”: [0, 0, 1, 0], “C”: [0, 0, 0, 1]). The length of the input vector was set at 400 bases. The sequences with length less than 400 nucleotides were padded for their empty columns by zero. Since each base was converted into a four-dimensional vector, each sequence was vectorized into a matrix with a dimension of 400X4 (Input sequence length X Alphabets of the input). We also used the structural information of these RNA sequences which hold special importance in case of lncRNAs where structural conservation is more expected than sequence conservation. This information of structure for each sequence was obtained in the form of Dot-Bracket representation of the secondary structure of the RNA by using RNAfold of ViennaRNA Package v2.4.18 [29]. We encoded the structure information for each base’s corresponding dot-bracket assignment in the sequence with “one-hot” encoding (“(”: [1, 0, 0], “.”: [0, 1, 0], “)”: [0, 0, 1]) with a vector dimension of 400X3 (Input sequence length X Alphabet of Dot-Bracket representation). Here also, padding was done for the shorter input sequences. The one-hot encoded bases and structural information are thus independent of one another. Applying both separately in a bi-modal manner would map the structural information to the sequence information while building the network models. The one-hot encoded metrics were then used as the input for the multi model convolution neural network to train and build the models for lncRNAs identification.

To evaluate the performance of the CNNs, various number of hidden layers were tested and finally two hidden layers were selected in fully connected manner. The number of the nodes across both the dense hidden layers were tuned based on the number of filters used in the convolution layer. All the component layers were optimized for their suitable numbers and component nodes number by iterative additions. Similarly, the kernel size and strides were optimized by trying different values in incremental order. An exponential activation function based single node classification layer was used with Mean Absolute Error loss function to calculate the loss. “Adam” optimizer was used at this point to adjust the weights and learning rates. Adam is an improved stochastic gradient optimizer which automatically adjusts the learning rates, momentum, and regularizers, making it more efficient algorithm. Dropout layer was also used to reduce overfitting in the model. The batch size was set to 100 and the number of epochs was set to 50. The DeepPlnc was implemented in Keras using TensorFlow. All codes were written for Python3. All the hyper-parametric values in the combination were tested and fixed using an in-house developed script to get the best hyperparameter values combination. This entire optimization process was done using two different approaches: Random search optimization and Bayesian optimizations. Full details of convergence to the optimum performance with hyper-parameters optimization is given in Supplementary Table S2, and Supplementary Figures S1, S2, and S3. The final layer outputs a confidence probability score indicated the confidence of each instance as non-lncRNA or lncRNA. If confidence probability score (Cp) ≥ 0.50 the meant that the corresponding input sequence was identified as lncRNA, else a non-lncRNA for the chunk. For the full length sequences, if the average of the confidence probability of all the sub-sequence chunks for the given sequence was >0.50, the sequence was considered as lncRNA, otherwise as a non-lncRNA.

### 2.3. Performance evaluation criteria

The performance of the developed model was evaluated on the test sets while applying the standard performance metrics. Confusion metrices containing correctly and incorrectly identified test set instances were built for the model. Sensitivity defines the portion of the positive instances which were correctly identified as positives whereas the specificity value describes the portion of negative instances which were correctly identified as negatives. Precision estimates the proportion of positives with respect to the total true and false positives. F1-score was also evaluated which measures the balance between precision and recall. Besides these metrics, Matthew’s Correlation Coefficient (MCC) was also considered. MCC is considered as the best metric to fathom the performance where score is equally influenced by all the four confusion matrix classes (true positives, false negatives, true negatives, and false positives) [24]. Better the MCC score, the more robust and balanced model with higher degree of performance consistency.

Performance measures were done using the following equations:

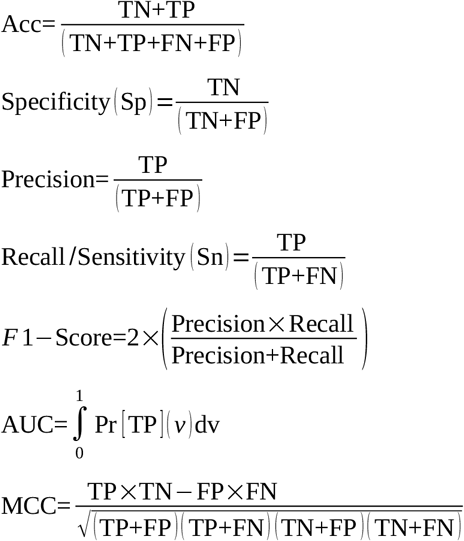

Where:

TP = True Positive,
TN= True Negatives,
FP = False Positives,
FN = False Negatives,
Acc = Accuracy,
AUC = Area Under Curve (AUC) from receiver operating characteristic (ROC) curves,

F1-score is a harmonic average of sensitivity and precision. MCC indicates a correlation coefficient between the true classes and the predicted classes.

### 2.4. Benchmarking and performance evaluation

To evaluate the DeepPlnc performance in this study, we compared DeepPlnc with five different tools: PLncPRO[13], CNIT [16], PreLnc [15], PlncRNA-Hdeep [19], and RNAplonc [14]. Two different datasets were considered separately for the benchmarking process as already described above (Dataset “A” and Dataset “B”). The four out of the compared five tools viz. PLncPRO, CNIT, PreLnc, and RNAplonc provide pre-built models. Only PlncRNA-HDeep does not provide any pre-built models. To overcome this problem, we built the model as described in its publication [19]. Both PLncPRO and PreLnc provide two different models for lncRNA classification. For PLncPRO, monocot and dicot are the two models available whereas for PreLnc, model built from *Zea mays* and *Arabidopsis thaliana* were available for prediction. In DeepPlnc, the sequences are scanned by overlapping windows of 400 bases at a time. For the identification with DeepPlnc, if more than 50% of the 400 bases long overlapping windows scored higher value for lncRNA (Cp ≥ 0.50) then the sequence was labeled as lncRNA, otherwise non-lncRNA. The sequence having length < 200 bases were discarded as they did not fit one of the formally accepted prime criteria to define lncRNAs. Both the considered datasets were tested in the form of full length complete sequences sets as well as randomly chopped sequences to mimic the incomplete sequences.

Besides these benchmarking tests, one more set of benchmarking tests was done where DeepPlnc was trained and tested across the datasets of the compared tools and the obtained performance was compared with the performance values claimed the respective tool.

## 3. Results and Discussion

### 3.1. Data collection and formation of datasets

A total of 9,911 experimentally validated plant ncRNAs sequences were downloaded from Ensembl Plants v51 [20], PncStress [21], and PLncDB V2.0 [22] databases for 51 plant species. Non-lncRNAs (mRNAs) data were downloaded from Ensembl Plants (v51) database for 51 plant species. The ncRNAs were used as positive data and the protein-coding transcript were used as negative data. This data contained 9,911 experimentally validated non-coding sequences. Sequences shorter than 200 nucleotides were discarded, keeping in the view that the minimum length of 200 bases is considered for lncRNAs. After filtration, this data got reduced to 6,563 sequences for 30 species. For non-lncRNA transcripts, 6,563 sequences were randomly obtained for the corresponding species considered for the lncRNA data. This dataset became the foundation data for training the deep learning modules. Initially, the above data was broken into two halves for training and testing datasets creation, containing 3,563 sequences of non-lncRNAs and 3,563 sequences of lncRNAs for training. Remaining 3,000 non-lncRNAs and 3,000 lncRNAs sequences were used purely as the test samples. In this entire study this dataset is referred as Dataset “A”. Supplementary Table S3 depicts the distribution of collected data across the various species in plants for Dataset “A”. This also needs to be mentioned here that this entire dataset split into training and testing sets has been done 10 times for 10 fold independent random trials for validation, building a new model every time from the scratch and every time absolutely no overlap of instances between training and testing data, ensuring no training memory in order to get the unbiased performance measure. This step was necessary to estimate the performance and error consistency of the model building approach.

Dataset “A” was also a neutral dataset where experimentally validated cases were considered and which was not used by any other tool considered here for comparison for model building. However, in order to provide an extra layer of benchmarking, one more dataset was made. This dataset was built using the data sources like PNRD database [23] and test datasets used in CNIT tool which were not used by the compared tools for their datasets and model building but followed the similar approach of annotating and classifying the lncRNAs. A total of 6,621 plant lncRNAs were obtained from PNRD database for 14 plant species and 439 lncRNAs were collected from the test set considered by CNIT for two plant species. A total of 7,060 lncRNA transcripts having minimum sequence length of 200 bases for 16 plant species were considered as the positive dataset. Non-lncRNAs data were downloaded from Ensembl Plants (v51) database for the corresponding 16 species considered for the lncRNA data. To generate a balanced test set, 7,060 non-lncRNA sequences were randomly selected from non-lncRNAs dataset for each species which accounted for the negative dataset. In the lncRNA dataset (positive), majority of the lncRNAs (93.8%) were predicted by various computational methods where all RNAs were compared with the coding RNAs in RefSeq and Ensembl and later using CNIT these RNAs were filtered. Only the filtered RNAs were kept and considered as noncoding RNAs. For the rest of the sequences, 6.2% of the total (7,060) for two plant species were reported as “known” sequences which were extracted from CNIT website (http://cnit.noncode.org/CNIT/) and did not overlap with Dataset “A” because it did not belong to the plant species which were considered for the Dataset “A”. This way another neutral dataset was created using lncRNAs and non-lncRNAs, referred as Dataset “B” in this entire study. Supplementary Table S3 depicts the distribution of collected data across the various species in plants for Dataset “B”. This dataset was not applied for training purpose and was used exclusively for testing only. Figure 1 provides the illustration about the creations of these datasets.

### 3.2. Parameter optimization and establishment of the final bi-modal CNN architecture

Figure 2 provides the details of the evolved CNN architecture to characterize the lncRNAs in plants. The framework of the bi-modal CNNs used a 2D CNN layer, one max-pooling layer, and three fully connected layer with one output layer along with two batch normalization layers to control the over-fitting during the training process. We used bi-modal CNNs with two different parts which used data encoding by one hot encoding method into binary matrix as shown in Figure 2. In the first part of the bi-modal CNNs (left side of Figure 2a) the nucleotide sequence worked as the input. The input layer was followed by a convolution layer containing 32 channels at each position with 3×3 kernel size with sequence dimension of 400×4×1 (Input sequence length X alphabets of the input X Input’s depth). The input sequence was padded if the length was shorter than 400 bases in order to ensure a constant size of the input matrix. The output resulted into 400×4×32 dimension representation after convolution. This output was normalized by the second layer called batch normalization layer. This layer was applied with parameters like momentum = 0.99 and epsilon = 0.001 to overcome the over-fitting problem during training process. This layer was followed by a third layer called max-pooling layer. This layer included 32 nodes having kernel size 2×2 with input dimension 400×4×32. Max-pooling helped in reducing the dimensions of convoluted sequence into a dimension of 200×2×32 where the stride size was S=2. Next to this layer, pooled feature maps were passed to fully connected layer by flattening it into one dimensional form having an input vector of 12,800 nodes. The hidden state layers in the present study had two dense layers with both having 32 hidden nodes with RELU activation function. The hidden layer output came in the form of 32 nodes dimension which was concatenated along with the output from the second CNN module as described below.

**Figure 2:**
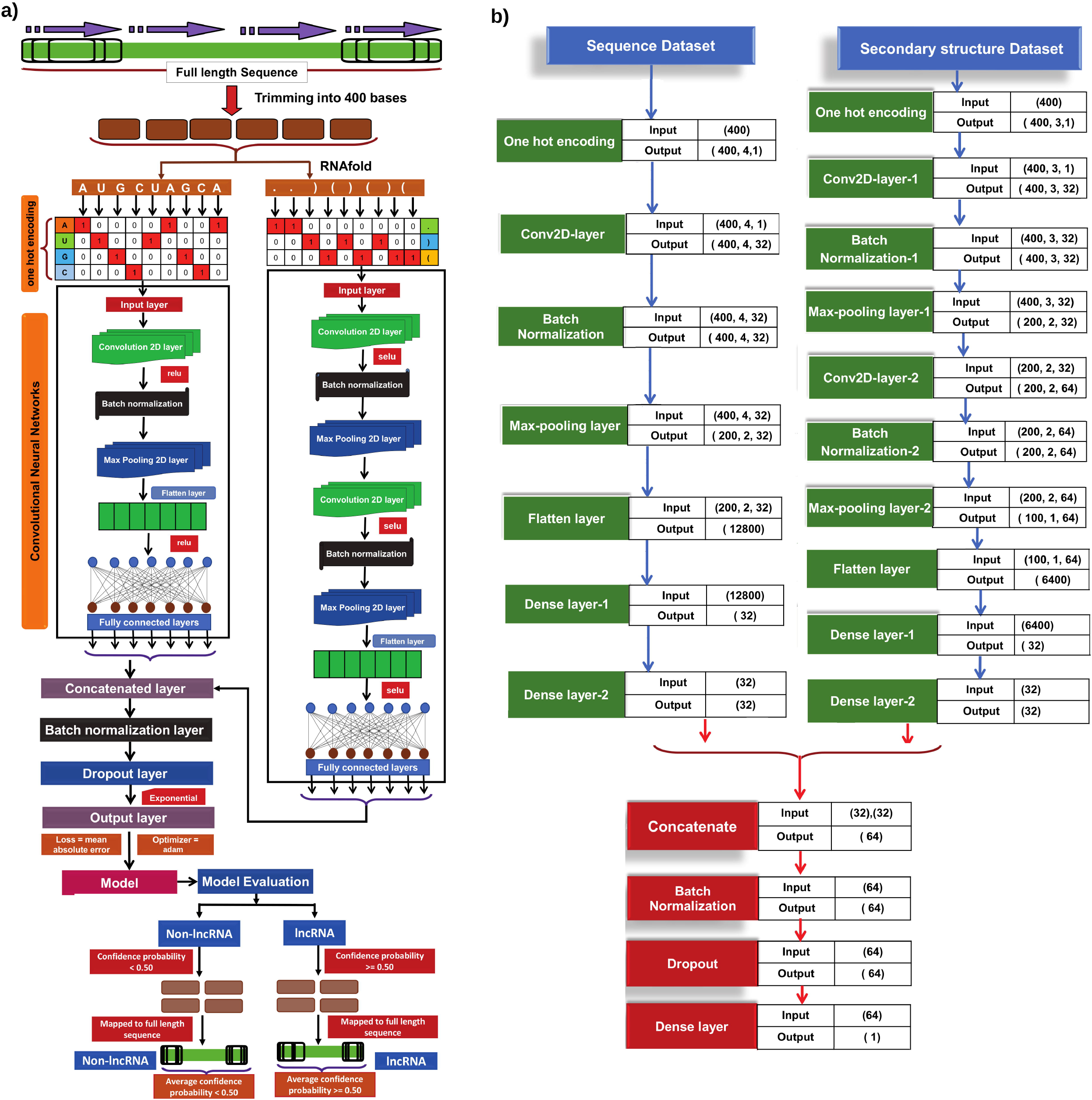
Detailed pipeline of the workflow. **a)** The image provides the brief outline of entire computational protocol implemented to convert full length sequences into 400 nucleotides length sequences that were further processed for feature generation such as nucleotide sequence and secondary structure derived from RNAfold tool for each block respectively and used as input features for bi-modal convolution neural network. **b)** Architecture and visualization of bi-modal CNN having two blocks, one Convolution Neural Network and second fully connected layers with input and output dimensions of each layers.

A similar CNN architecture was followed by the second part (the right side of Figure 2a) of the bi-modal CNNs which considered secondary structural information in the form of dot bracket representation (“(“, “.”,”)”) for the input sequence. The input layer passed on the input to the convolution layer having a dimension of 400×3×1 (Input sequence length X Alphabets in Dot-Bracket secondary structure representation X Input’s depth) with 32 channels, 3×3 kernel sizes, stride size of one, and along with SELU activation functions with kernel constraints having maximum value 3. Inputs were padded with zero to ensure a constant size of the matrix in case of shorter input sequences. The second layer was max pooling which used normalized instances by passing through a batch normalization layer with input shape of 400×3×32 with stride size of two, padded with zero for a constant matrix size. That gave output in the form of reduced features maps with dimension of 200×2×32. This output was passed to the second convolution layer having a dimension of 200×2×1 with 64 channels, 3×3 kernel sizes, stride size of one, and along with SELU activation functions with kernel constraints having maximum value of “3”. Inputs were padded with zeros to ensure a constant size of the matrix in case of shorter input sequences. The fourth layer was max pooling which used normalized instances by passing through a batch normalization layer with input shape of 200×2×64 with stride size of two. This gave reduced features map output with 100×1×64 dimension. This output became the input to a system of two hidden layers with fully connected nodes with 32 nodes per hidden layer with SELU activation function. It used an input size of 6,400 nodes.

At the end, the output from these two CNNs based networks were connected into one concatenated layer that received input in the form of 32+32 nodes, returning a vector of 64 elements. This vector was processed by one batch normalization layer and was passed into a dropout layer with dropout fraction 0.2. By passing 0.2 dropout fraction, 20% of the hidden units were randomly dropped during the training process of the model. This layer helps to reduce overfitting. It ensured almost negligible difference between the training and validation performances. Finally, the output of the dropout layer with 64 dimension was passed to the last and final output layer, a node with exponential activation function. The model was compiled by mean absolute error loss function to calculate the loss which was optimized by using the “Adam” optimizer. The output of the last layer (output layer) returned the confidence probability for each input sequence while working on a window size of 400 bases (chunks). The confidence score indicated the confidence of each instance as non-lncRNA or lncRNA. Based on this confidence probability score (Cp), if the Cp is ≥ 0.50 means the corresponding input sequence was identified as lncRNA and if the Cp is < 0.50 then it is non-lncRNA for the chunk. For the full length sequences, if the average of the confidence probability of all the chunks for a particular sequences was found ≥ 0.50, the sequence was considered as lncRNA, otherwise as a non-lncRNA. The final hyperparameters set for the output layer of the implemented model was: {“Activation function”: Exponential, “Loss function”: mean absolute error, “Optimizer”: Adam}. The related information about optimization towards the final model is listed in Supplementary Table S2 and illustrated in Supplementary Figures S1, S2, and S3.

### 3.3. Ablation test for the properties effect on the network and performance assessment

For the determination of the optimal feature combinations in the lncRNA classification model, we used six properties combinations for various input lengths for different input property encodings: 1) Monomers density, 2) Secondary structure, 3) Monomers density + Secondary-structure, 4) Dinucleotides density + Trinucleotides density, 5) Monomers density + Secondary-structure + Dinucleotides density + Trinucleotides density, 6) Monomers density + Secondary-structure + number of hydrogen bonds + purine-pyrimidine sequence encodings. Initially, the considered dataset was split into 50:50 ratio to form the training and testing datasets. Then, the model was trained with the training set sequences and all the above mentioned properties encodings were done accordingly for that window size in sliding window manner. On the test set for the above mentioned input property encoding combinations, the accuracy values scored were 83.64%, 71.34%, 92.54%, 77.63%, 87.54%, and 89.48%, respectively. The window length was varied from the length of 200 to 800 bases and the scorings were obtained for all these window lengths. Best performance was observed at window size of 400 bases. Even the padded shorter sequences with shorter lengths as low as 200 bases performed the best for this window size. For the chunked dataset, the average accuracy values scored for this window based classifiers were 85.17%, 68.68%, 98.15%, 87.54%, 95.64%, and 93.54%, respectively.

Figure 3(a) provides a snapshot of impact of the variable window sizes and combinations of property encodings considered for this chunked dataset. Finally, the combination of monomeric and dot-bracket secondary structure representations with a sliding window of 400 bases was used for the final model training which scored an initial accuracy of 98.15% on the chunked dataset (98.99% sensitivity, 97.31% specificity, 98.17 F1-score, 0.96 MCC score). To take the final decision on the full length input sequence, all chunks were mapped back to the original input sequence and were assigned to the class lncRNA if average score of all the chunks was found higher than 0.5. In this way, the average accuracy of the above classifier for the complete length sequence increased and attained the accuracy value of 98.78% with 99.37% sensitivity, 98.20% specificity, 98.79% F1-score, 0.97 MCC score.

**Figure 3:**
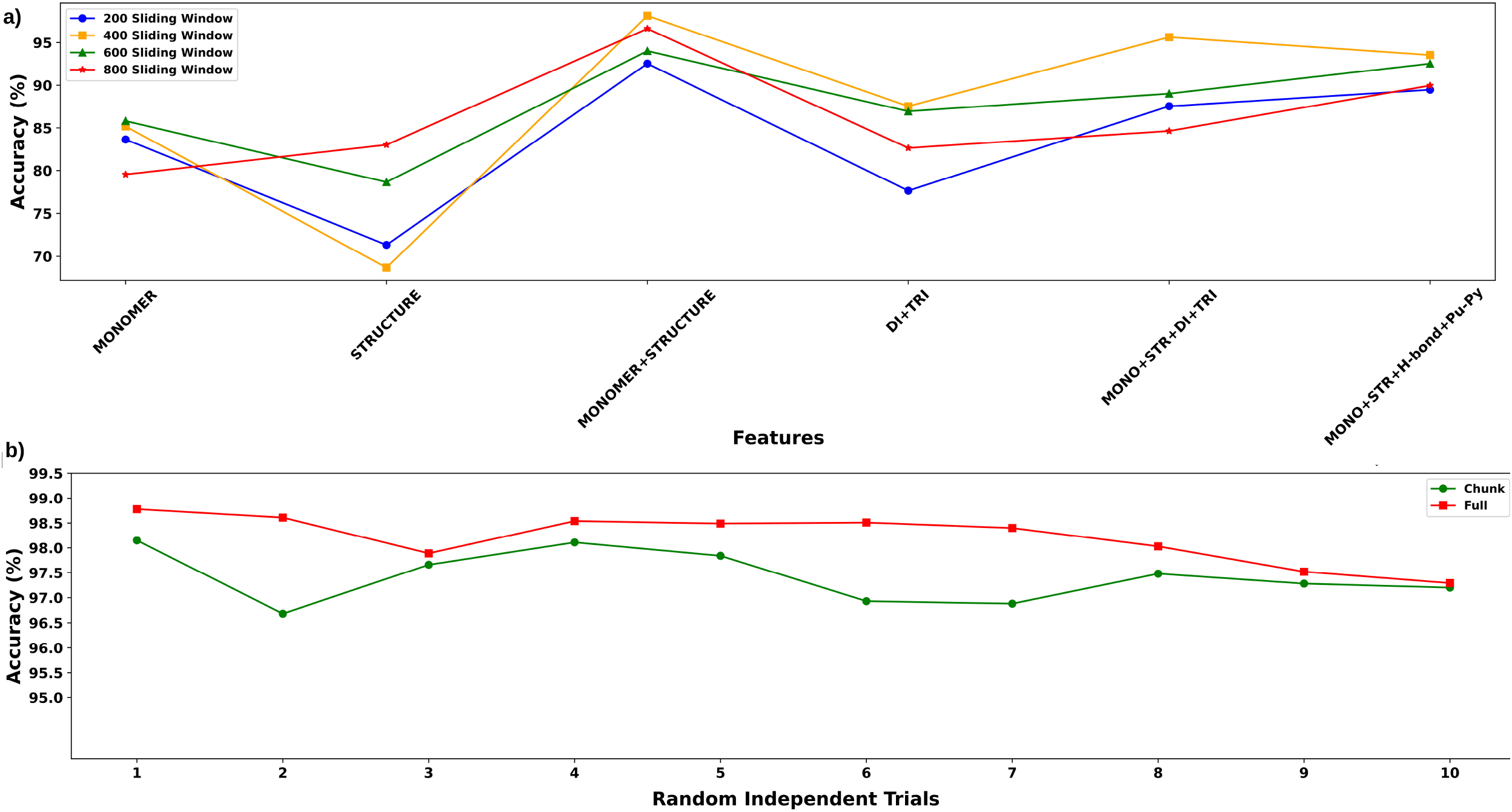
**a)** Accuracy plot for different feature combinations used to select important features. Third combination which was monomer + secondary structure reported higher accuracy than other combinations [monomers, Structure (Secondary structure), Dinucleotides, Trinucleotides, H-bond (Hydrogen bond), Pu-py (Purine-Pyrimidine)]. **b)** Ten-fold independent random trials for validation of DeepPlnc on Dataset “A” depicts consistent performance of the tool on randomly shuffled data to train and evaluate the model.

The next step was 10 fold independent random trials with 10 different pairs of randomly formed train and test sets in order to verify the consistency of the performance of the model (Supplementary Table S4). 10 times from the dataset, a totally new model was built from the scratch while using randomly selected pairs of train and test sets every time, and the performance was noted. Here also, no overlap existed between the train and test sets in order to ensure no bias and memory. As can be found from the Figure 3(b), the developed system performed consistently good and scored high on all these training and testing combinations where the average accuracy consistently scored above 97% and breached 98% for the complete length input sequences. Also, not much fluctuations in performance was observed and consistently performed around 98% accuracy, underling the robustness and consistency of the approach. As transpires from the scoring obtained for the various metrics, the model performed consistent and well balanced, besides scoring high accuracy across all the test data sets which were having experimentally validated cases of non-lncRNAs and lncRNAs.

### 3.4. Benchmarking: DeepPlnc consistently outperformed all the compared tools

A highly comprehensive benchmarking study was performed where DeepPlnc was compared with five different recently published tools to detect plant lncRNAs: PlncPRO, CNIT, PreLnc, PlncRNA-HDeep, and RNAplonc. Besides this, the benchmarking also considered two different datasets (“A”, “B”; already detailed above) to carry out a totally unbiased assessment of performance of these tools on different datasets. Each dataset was further divided into two sub-categories, chunked form containing incomplete RNA sequences of size 400 bases (shorter sequences were padded) and full length sequences. This is important to note that the chunked data also provided a frame to mimic the incomplete/truncated length sequences which usually appear in large number in the *de novo* assembled transcriptome data as well as provide a realistic treatment to deal with those conditions where boundaries of lncRNAs are not known, like genomic annotations.

All these six software were tested across the two datasets (“A” and “B”) where DeepPlnc consistently outperformed all the compared tools for all these datasets, for both of their forms (incomplete chunked sequences and complete sequences), and for all of the performance metrics considered. This must be noted the test set from Dataset “A” was never seen before by the DeepPlnc as it was kept aside exclusively for the validation purpose only. The Dataset B worked exclusively as a totally validation test set as no part of it was used even for training purpose for DeepPlnc. This way it was ensured that both the test sets were totally unseen and fit for unbiased assessment.

Figure 4 gives a detailed view of the performance and benchmarking analysis across the two datasets studied for all these software. On the chunked transcript data representing the incomplete sequences, DeepPlnc scored 98.15% accuracy and 0.96 MCC scores, for Dataset “A”, taking a lead of 10.29% in accuracy and ~0.19 lead in MCC scores from the second best performing tool, PLncPRO. On the complete length sequences, DeepPlnc scored the accuracy of 98.78% and the MCC value of 0.97 on Dataset “A”, taking a lead of ~6.18% for accuracy and ~0.12 lead for MCC from the second best performer while emerging as the best performer by large margin. On the chunked dataset CNIT and PreLnc achieved an accuracy of 87.07% and 82.59% and MCC of 0.74 and 0.66, respectively. For the complete sequences they achieved an accuracy of 87.95% and 92.60%, respectively. Their MCC values were 0.76 and 0.85, respectively. PlncRNA-HDeep and RNAplonc were the least performing tools on the chunked dataset, scoring an accuracy of 65.66% and 61.35%, respectively. Their MCC values were 0.36 and 0.36, respectively. However, RNAplonc achieved an accuracy of 92.20% and MCC of 0.84 on complete sequence dataset whereas PlncRNA-HDeep achieves an accuracy of 74.62% and MCC of 0.51. Figure 4 (a) and (b) provide the complete performance benchmarking for this Dataset “A” for chunked incomplete sequences and complete lengthen instances, respectively.

**Figure 4:**
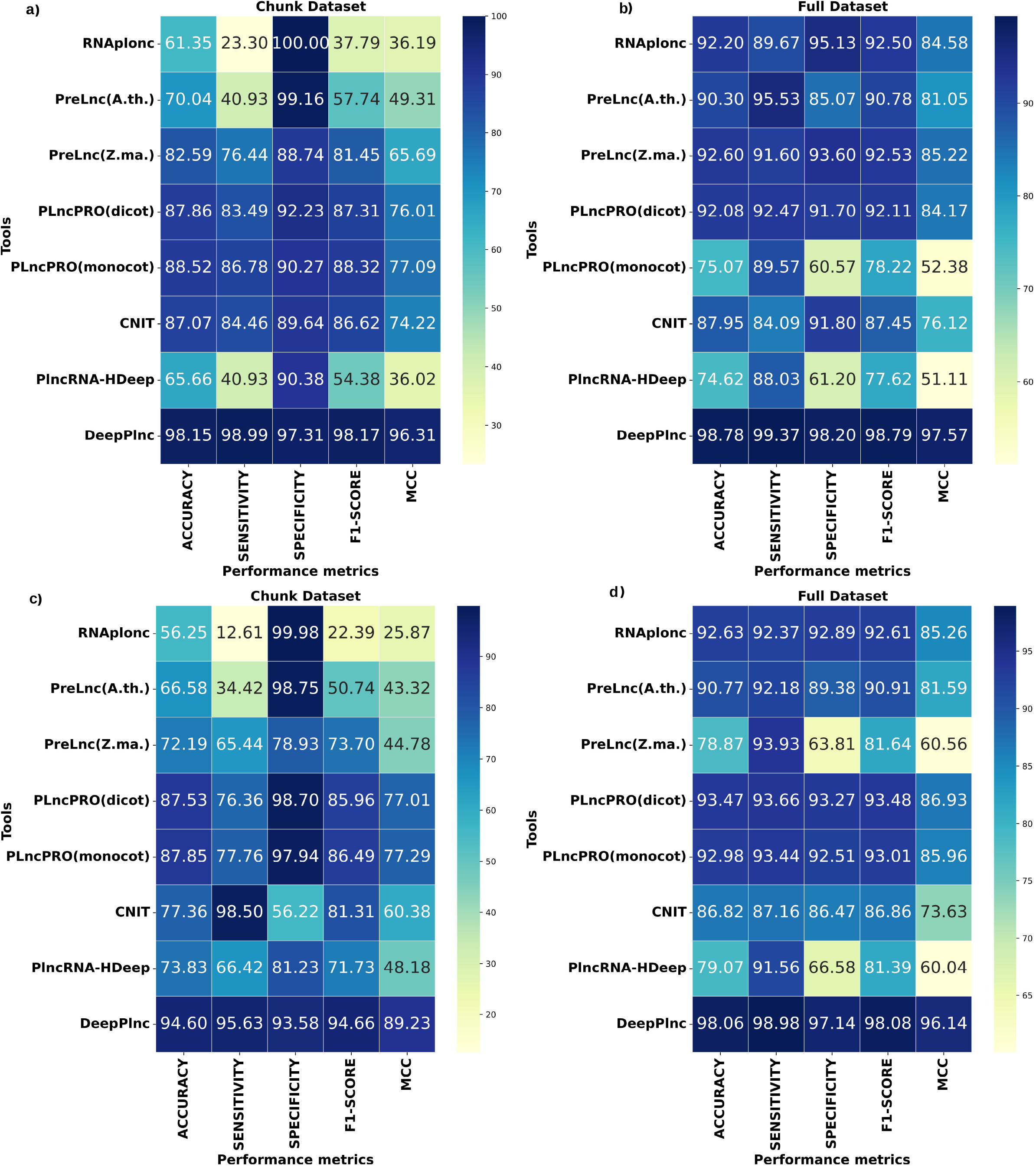
Comparative benchmarking with different combinations of test datasets. **(a)** Chunk Dataset “A”, **(b)** Full-length Dataset “A”, **(c)** Chunk Dataset “B”, and **(d)** Full-length Dataset “B”. For every such combinations, the performance metrics (sensitivity, specifcity, F1-score, MCC and accuracy) are given in the form of heatmap. The plots clearly indicate that DeepPlnc consistently outperforms in comparison to existing tools for all the metrics based on all these different combinations on different datasets. Consistently high MCC scoring obtained by DeepPlnc reflects it as a robust tool where dispersion in performance metrics was least and DeepPlnc also scored high sensitivity for all set of combinations, clearly depicts that it performs better than other plant lncRNA classifier tool. MCC values were converted to percentage representation for scaling purpose.

Coming to the Dataset “B”, DeepPlnc continued its top performance even on this dataset while scoring the highest values for all the performance metrics considered. It clocked an accuracy of 98.06% and the MCC value of 0.96 for this dataset on the complete length sequences. It scored 4.59% better accuracy than the second best performing tool, PLncPRO, and scored 0.1 higher MCC value than PLncPRO on the complete length sequences. On the truncated chunked data of Dataset “B” also, DeepPlnc continued to outperform others with good margin (Accuracy: 94.60%, MCC: 0.89) and maintained a margin of 6.75% better accuracy than the second best performing tool, PLncPRO. It also scored 0.12 better MCC score than PLncPRO. For the chunked dataset, CNIT and PreLnc achieved the accuracy scores of 77.36% and 72.19% and MCC values of 0.60 and 0.44, respectively. For the complete sequences they achieved an accuracy of 86.82% and 90.77% and MCC values of 0.73 and 0.81, respectively. PlncRNA-HDeep and RNAplonc were the least performing tools on the chunked dataset, scoring an accuracy of 73.83% and 56.25%, respectively. They scored MCC values of 0.48 and 0.26, respectively. However, RNAplonc achieved an accuracy of 92.63% and MCC 0.85 on the complete sequences. Whereas PlncRNA-HDeep achieved an accuracy of 79.07% and MCC value of 0.60. Figure 4 (c and d) provides the complete performance benchmarking for this Dataset “B” for the chunked incomplete sequences and complete length instances, respectively.

DeepPlnc scored an AUC value of 99.98% (incomplete sequences) and 99.69% (complete sequences) on Dataset “A”. For Dataset “B”, it scored the AUC values of 98.97% (incomplete chunked sequences) and 99.55% (complete sequences). As illustrated in Figure 5, after DeepPlnc, PLncPRO and PreLnc emerged as the next best AUC scorers. What is also noteworthy besides the high accuracy and F1-score values of DeepPlnc across all the datasets, is the fact that its MCC values always stood out as the highest. DeepPlnc maintained least variability and dispersion of performance scores and continued to display its strong balance in detecting the positive and negative instances with high and similar level of precision. A higher MCC value is considered as the true reflector of the robustness, consistency, balance, and performance as it becomes higher only when all the four metrics of performance measure score high [24].The full details and data for the benchmarking studies are given in Supplementary Table S5. Having such consistent performance lead where accuracy lead ranged from 4.6 to 10% from the second best performing tool is itself is a big margin. The observed performance lead of DeepPlnc was evaluated for significance using Chi Square test on the performance contingency tables having True Positives, False Positives, True Negatives and False Negatives for the compared tools. For all the comparisons, highly significant p-values were observed (p-value << 0.01), clearly suggesting that the observed performance lead of DeepPlnc was significantly higher than the compared tools. And when it comes to application in the genomic setup, such margin in performance becomes much bigger and significant where a margin of just 1% itself may cause several false cases.

**Figure 5:**
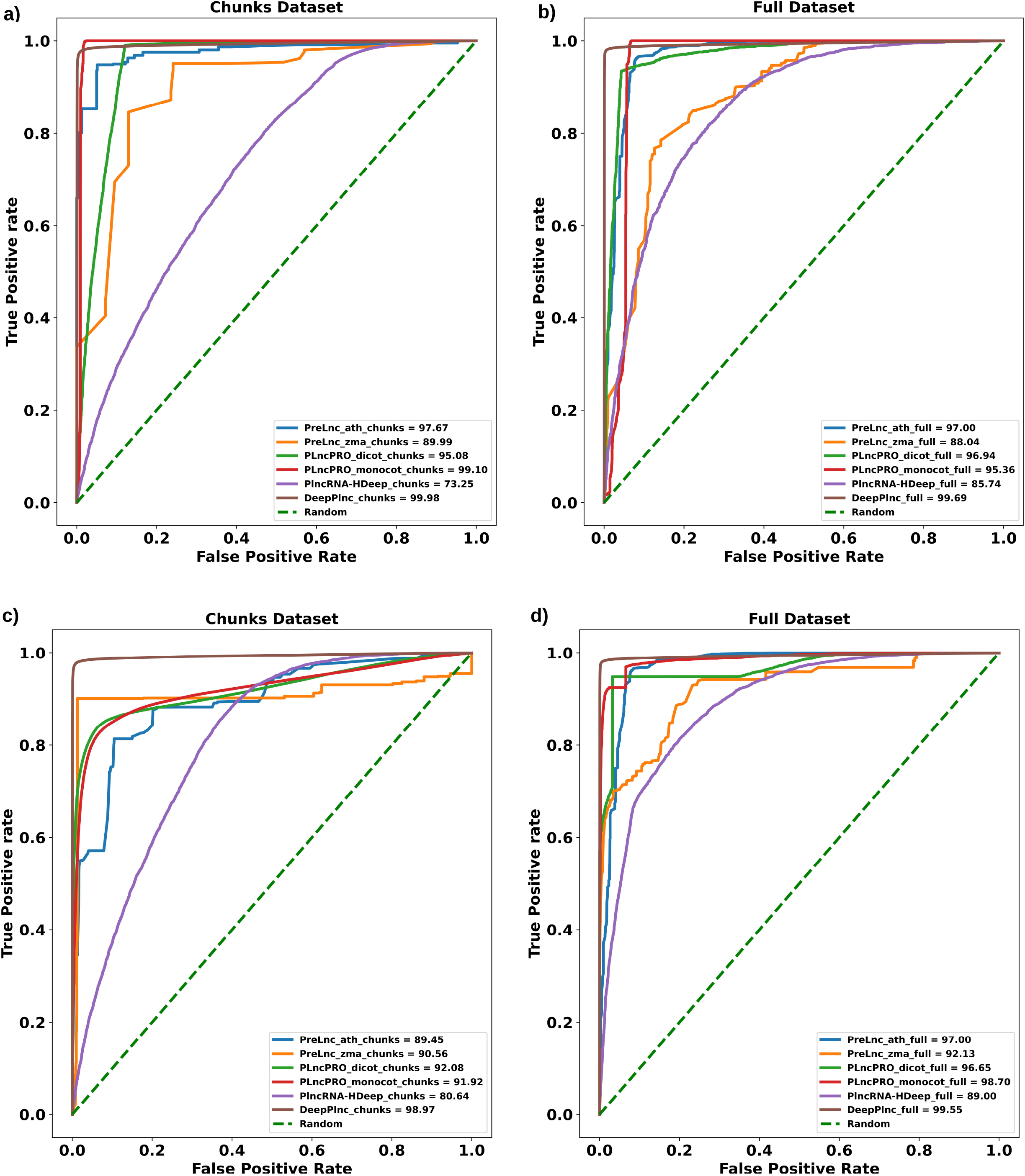
ROC curve plot obtained by DeepPlnc and all others tools for all combinations, **(a)** Chunk Dataset “A”, **(b)** Full-length Dataset “A”, **(c)** Chunk Dataset “B”, and **(d)** Full length Dataset “B”.

Besides the above mentioned comprehensive benchmarking tests, one more benchmarking test was performed. In this one-to-one comparison, DeepPlnc used the compared tool’s dataset to build its models from their train sets and tested its performance on their test sets. This way five times DeepPlnc models were built from the scratch and tested for the respectively compared tools. The observed performance of DeepPlnc was compared with the claimed performance of the compared tools, and for almost all the compared tools, DeepPlnc’s performance was superior (Supplementary Figure S4). It was also noted that all these compared software had far bigger gap between sensitivity and specificity values, suggesting even better balanced performance by DeepPlnc. Only PreLnc’s claimed accuracy was found slightly higher than DeepPlnc. PreLnc scored just 0.74% higher accuracy than DeepPlnc, but also displayed huge difference of 7.6% between its sensitivity and specificity values, suggesting imbalanced performance by PreLnc. In our previous benchmarking with both datasets, A and B, PreLnc had not even figured in the top three tools in terms of performance. And on the chunked incomplete sequences, its performance was among the bottom two. Thus, it was needed to verify the performance of PreLnc on the datasets used by other tools and compare its performance there with DeepPlnc’s performance on the same datasets. PreLnc was run across the datasets of all the remaining compared tools and its performance was compared with DeepPlnc’s performance for the same datasets and compared tools. PreLnc was found outperformed by DeepPlnc across all these four tool’s datasets (Supplementary Figure S4 (C)), besides being outperformed during bechnmarking with Datasets A and B.

This way, through a very comprehensive benchmarking across a wide range of datasets, it was established that DeepPlnc consistently performed as well as outperformed all the compared tools.

### 3.5. Application demonstration: Detection and annotation of novel lncRNAs in *Rheum australe* transcriptome data

To demonstrate the applicability of DeepPlnc, it was run against the recently published transcriptome data of an endangered Himalayan medicinal plant, *Rheum australe* whose genome sequence is still not known and absolutely no information is there for lncRNAs [25]. The assembled transcriptome of *Rheum australe* had reported a total of 35,679 transcriptome sequences, of which 23,569 sequences were having significant BLAST hits. As mentioned by Mala et al., 2021 [25], dissimilar sequence (DS) clustering was performed to identify the multiple representatives of the same gene which reduced the number of total unigenes from 23,569 to 21,303. These 21,303 unigene transcripts comprised well annotated protein coding RNAs (3,912), hypothetical (7,109) and predicted genes (10,282). Besides them, without any significant BLAST hit, a total of 12,110 transcripts stood as the potential candidates for lncRNA annotation. In this study, before going into further analysis, for the sake of clarity we removed the predicted and hypothetical protein coding sequences form the study because there was no experimental evidence that if such sequences really coded proteins or not and fully annotated protein coding sequences. The well annotated non-lncRNAs were kept knowingly as its characterization would provide the extent of error done by the software on the false positive front. A total of 2,991 out of 3,912 non-lncRNAs with minimum length of 200 bases were selected to annotate the non-lncRNAs whereas 921 sequences were discarded due to shorter length. In the similar way, a total of 6,598 transcripts were taken from 12,110 transcripts as input to DeepPlnc with minimum length of 200 bases to annotate the lncRNAs and rest of the 5,512 sequences were discarded. Figure 6 depicts the flow diagram representation of the process of lncRNA identification in *Rheum australe* and the result at its every step of execution.

**Figure 6:**
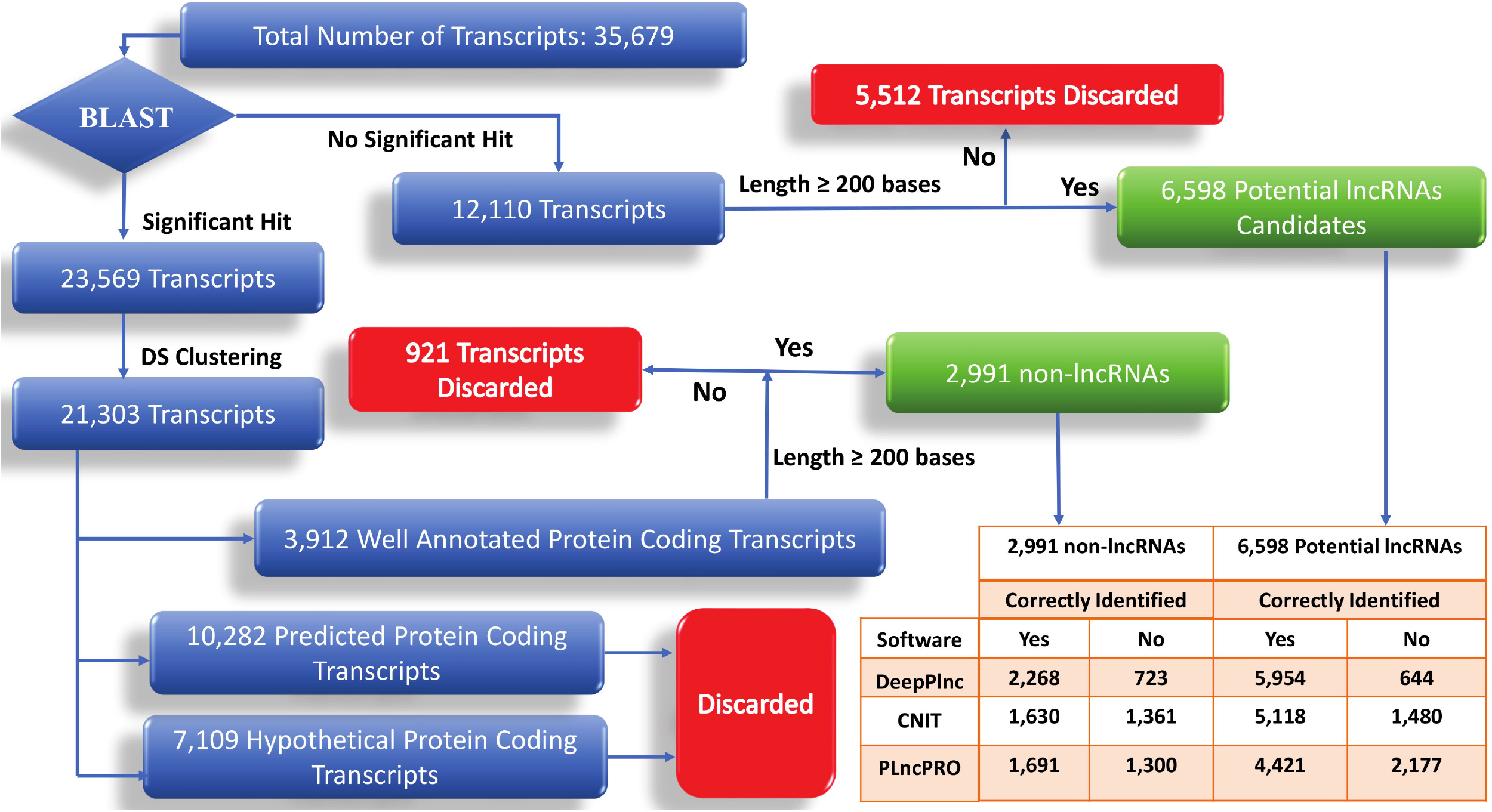
Flow diagram representing the process of lncRNAs identification in *Rheum australe.* Compared to other tools, performance of DeepPlnc was found much superior. Its wrong classification for the well annotated RNAs as lncRNAs was the least. Also, it classified most of the potential lncRNAs correctly where also it performed better than others.

After this filtering process we had 2,991 transcripts as non-lncRNAs candidate and 6,598 transcript as potential lncRNAs candidate. A total of 5,954 transcripts out of 6,598 potential lncRNAs candidate were identified as lncRNAs and 644 transcript were identified as non-lncRNAs by DeepPlnc. Similarly, a total of 2,268 transcript were identified as non-lncRNAs out of 2,991 non-lncRNA candidate transcripts whereas 723 transcripts were wrongly classified by DeepPlnc. However, ~90% of the lncRNA transcripts were classified as the lncRNAs. As implemented in DeepPlnc, sequences having average confidence probability ≥0.50 were classified as lncRNAs and if the average confidence probability was < 0.50 the sequence was classified as a non-lncRNA. In order to evaluate how other lncRNA identification tools characterized the data and how much wrong classifications they did to the well annotated non-lncRNA data, the two good performing tools from the benchmarking study described above, PLncPRO and CNIT, were also used with their default parameters to annotate this transcriptome data (Supplementary Table S6). CNIT classified 5,118 transcript as lncRNAs out of 6,598 potential lncRNAs and 1,480 transcripts as non-lncRNAs. Whereas, only 1,630 non-lncRNAs were correctly identified out of 2,991 non-lncRNAs candidate. Similarly, PLncPRO annotated 4,421 transcript as lncRNAs out of 6,598 potential lncRNAs and identified only 1,691 transcript as non-coding transcript out of 2,991 non-lncRNAs candidate. Figure 7 provides a Venn diagram of results obtained for this analysis and how these three tools performed on *Rheum australe* transcriptome annotation. A total of 6,677 and 2,912 transcripts were identified as lncRNA and non-lncRNA, respectively, through DeepPlnc. A total of 5,721 and 3,868 transcripts were identified as lncRNA and non-lncRNA, respectively, through PLncPRO. A total of 6,479 and 3,110 transcripts were identified as lncRNAs and non-lncRNAs, respectively, through CNIT. The last part of Figure 6 provides the overall result produced by all these tools, where it is very clear that DeepPlnc gives at least half the lesser false characterizations than the other two tools, which is a highly significant margin while annotating any genomic system. These values underlined the levels of confidence one could put on these tools when performing the annotation job. Also, it is possible that some of these transcripts annotated in the root study may not be really coding in actual scenario and some a noncoding pseudogene transcripts as many of them are supposedly incomplete sequences.

**Figure 7:**
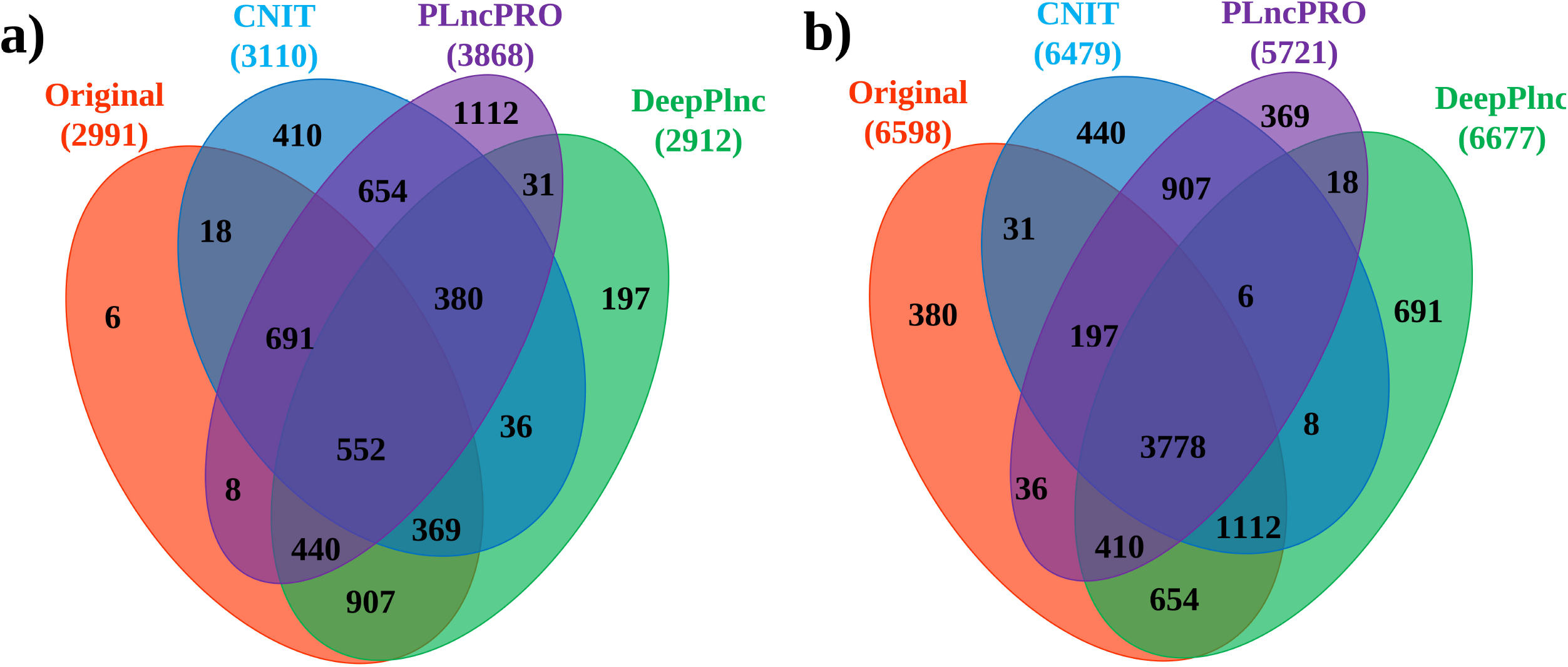
**a)** Venn diagram showing the number of coding and long non-coding transcripts annotated by CNIT, PLncPRO, and DeepPlnc on coding dataset. **b)** Venn diagram showing the number of coding and long non-coding transcripts annotated by CNIT, PLncPRO, and DeepPlnc on the long non-coding dataset. Common entities in two or more tools are enclosed in the overlapping portion of the ellipses.

Further delving on the transcriptome data was done for expression measures to assess how well the usual observation that lncRNAs are usually lesser expressed than non-lncRNAs. This was reflected by the identified lncRNAs in the transcripts of *Rheum australe*. The expression level which was measured in Fragments Per Kilobase of transcript per Million mapped reads (FPKM) of all the identified lncRNAs was compared with that of non-lncRNAs were calculated using RSEM [26] (Supplementary Table S7). The expression level of lncRNAs was found indeed lesser than non-lncRNAs at 4°C (t-test p-value 0.01). However, this gap in the expression was not that sharp at 25°C presumably due to lowering of the gene expression at higher temperature which is noticed in this Himalayan plant in response of higher temperature. The lncRNAs were found higher expressed at 4°C than 25°C (t-test p-value of 0.007). Since these tests were sensitive to some extreme values due to which mean values get influenced, medians/ranks based non-parametric ranking tests (Mann Whitney test) were also done. All these observation returned much higher significance, suggesting significant differences between lncRNAs expression and non-lncRNAs expression. Significantly higher expression of the lncRNAs was observed at the lower temperature for this plant. The mean and median expression values for lncRNAs at the lower temperature were 35.92 and 6.37, respectively. The same for non-lncRNAs were 42.67 and 8.94, respectively, at the lower temperature. At the higher temperature, the mean expression value was 16.51 and median expression value was 1.47 for the lncRNAs. While the non-lncRNAs displayed 20.36 mean expression value and 3.26 median expression value at the higher temperature.

For differential expression analysis, a total of 5,954 lncRNAs, as identified by DeepPlnc, were considered. In order to find the differentially expressed lncRNAs (DELs) at the two different temperatures (4°C and 25°C). A comparative analysis was performed for 4°C vs 25°C expression status. Significantly up- and down-regulated DELs were identified using edgeR [27] with log fold change (FC) ≥ 2 at a statistical significance level of p ≤ 0.05 and false discovery rate (FDR) ≤ 0.05. A total 2,420 DELs were identified out of 5,954 lncRNAs, of which 689 were up-regulated and 1,731 were down-regulated at the lower temperature as compared to that at higher temperature (Supplementary Table S8). At 25°C, at total of 1,731 lncRNAs were found over-expressed, while at 4°C, at total of 689 lncRNAs were found over-expressed. Supplementary Table S8 contains the list of such differentially expressed lncRNAs.

This all demonstrated the extent to which, tools like DeepPlnc could be important and useful in the annotation of totally new transcript datasets which contain a large number of incomplete sequences while giving much superior and reliable performance than previously published tools.

### 3.6. Webserver and standalone implementation

DeepPlnc has been made freely available a very simple to use webserver as well as a stand-alone program. The user needs to paste the RNA sequences in FASTA format into the text box or upload the RNA sequences in FASTA format and then click the submit button for the identification. This data is run through the trained bi-modal CNN which generates a relative probability scores for every position in the sequences. Later, the result page appears from where results can be downloaded in a tabular format and sequence wise distribution of probability score in the form of line plot as well as violin plot is displayed on the result page in an interactive form. The server has been implemented using D3JS visualization library, Python, JavaScript, PHP, and HTML 5. Figure 8 provides an overview of DeepPlnc webserver implementation. Besides this, a stand-alone version is also provided at GitHub at https://github.com/SCBB-LAB/DeepPlnc/.

**Figure 8:**
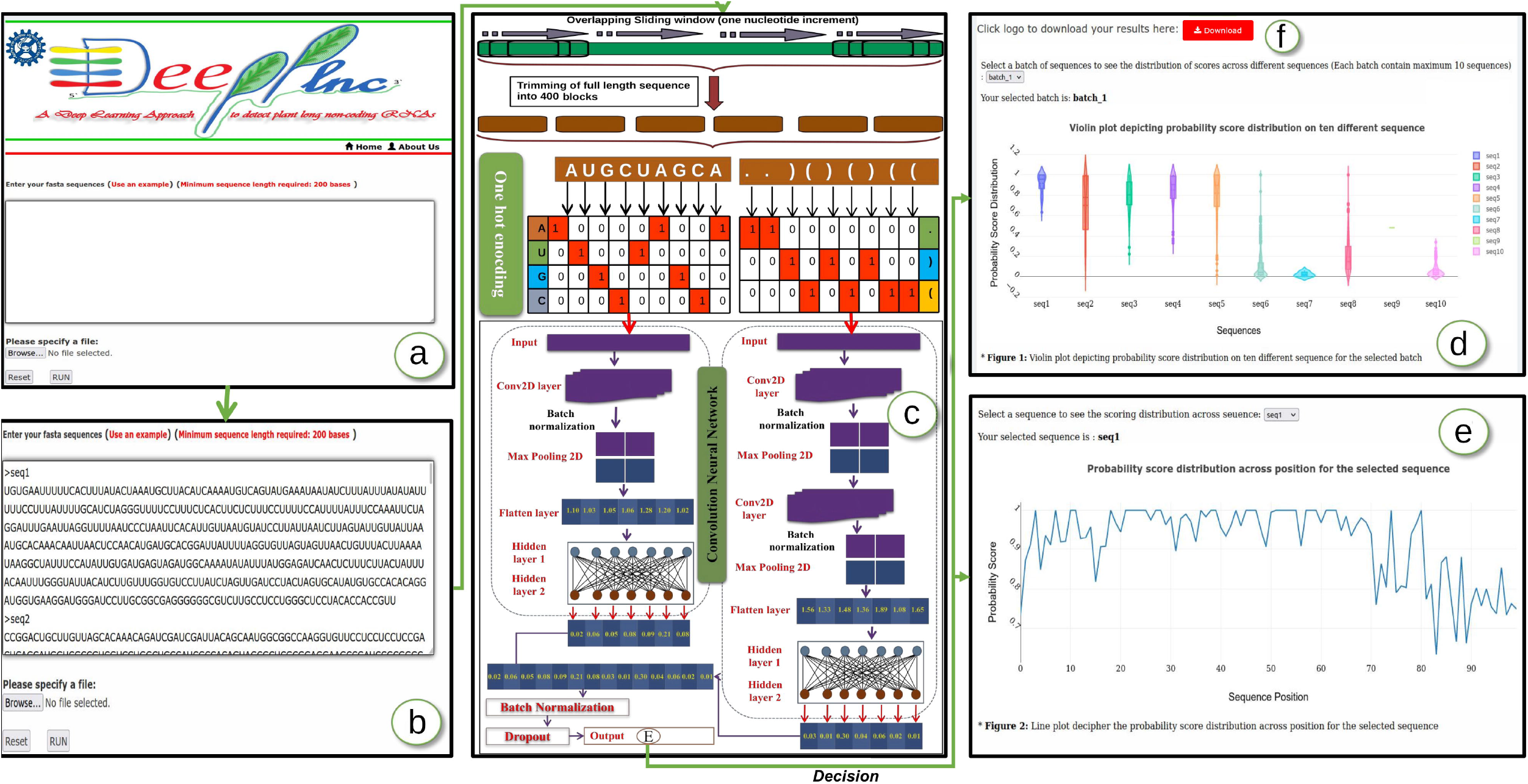
DeepPlnc webserver implementation. **a)** Input data box where the user can either paste or load the input file, **b)** Input is RNA sequence in FASTA format, **c)** These sequences were chopped into sequences of length 400 base pair in overlapping fashion and their secondary structure were obtained by running RNAfold. Later, sequence and their secondary structure represented by triplets were converted into one hot representation which was finally fed into the bi-modal CNN which in turn gives the position wise probability scores, **d)** These probability scores are represented in the form of interactive violin plot depicting the distribution of probability score on the ten different sequence for the selected batch, **e)** Also, the probability scores are represented in the form of interactive line plot depicting the distribution of probability score across position for the selected sequence. **f)** Download option for the result in the tabular format where the first column indicates sequence ID and second column represents whether it is lncRNA or not.

## 4. Conclusion

The lncRNAs have emerged among the important components of the eukaryotic transcriptomes and regulatory systems. A growing number of lncRNAs are now being found implicated in plant growth, development, and maintenance. There have been some tools developed for plants to identify lncRNAs but their inconsistency and performance gaps leave ample space to develop new approaches. Development with state-of-the-art approaches like deep learning remained an underexplored area in plant lncRNA biology. The present work provides one of the first deep learning approaches which returns high performance scores for all the considered metrics across a large volume of experimentally validated test datasets. It significantly outperformed most recent and prominent tools for plant lncRNA identification with high consistency and balance. DeepPlnc, was also found very accurately detecting even the incomplete transcripts. Such results make it a very important tool for annotating *de novo* transcriptome data which are studded with incompletely assembled transcripts and highly prone towards wrong annotations. Its capability to work on boundary-less inputs makes its practical application for highly reliable annotations of genomes and transcriptomes.

## Declarations

### Availability of data and materials

All the secondary data used in this study were publicly available and their due references and sources have been provided. All data and information generated/used, methodology related details etc. have also been made available in the Supplementary data files provided along with and also made available through the related open access server at https://scbb.ihbt.res.in/DeepPlnc/.

## Acknowledgements

The study was carried out under the aegis of The Himalayan Centre for High-throughput Computational Biology (HiCHiCoB), a BIC supported by the Dept. of Biotechnology, Govt. Of India. We are thankful to CSIR for the support through iPRESS project. Ritu is thankful to CSIR for her fellowship and to Academy of Scientific and Innovative Research (AcSIR) for her Ph.D. enrollment. SG is thankful to CSIR project iPRESS and DBT BIC support under HiCHiCoB for his project associateship. NKS is thankful to CSIR project iPRESS for his senior project associateship. All authors are thankful to the Director, CSIR-IHBT, for his kind support for this study. The IHBT MS communication ID is 5021.

## Authors’ contributions

Ritu and SG carried out the study. NKS assisted in the study. RS conceptualized, designed, analyzed and supervised the entire study. Ritu, SG, NKS, and RS wrote the MS.

## Supplementary information

**Supplementary Table S1:** Source information of Dataset “A” and Dataset “B” used to construct, evaluate and benchmark the DeepPlnc bi-modal CNN.

**Supplementary Table S2:** Hyperparameter optimization for DeepPlnc.

**Supplementary Table S3:** Number of experimentally validated, known and predicted lncRNAs used to construct, evaluate and benchmark DeepPlnc for Dataset “A” and Dataset “B”.

**Supplementary Table S4:** Tenfold independent random trials for validation of trained bi-modal CNN model.

**Supplementary Table S5:** Benchmark performance of DeepPlnc and other selected tools on two diiferent datasets.

**Supplementary Table S6:** Classification performance of tools on *Rheum australe* transcriptome. **Supplementary Table S7:** FPKM values of lncRNAs detected by DeepPlnc in *Rheum australe.* **Supplementary Table S8:** Diffrentially expressed lncRNAs at 4 °C vs 25 °C.

**Supplementary Figure S1:** Optimization results for hyperparameters for sequence part of the bimodal CNN. **A)** Batch size optimization, **B)** Dropout rate optimization, **C)** Epoch size optimization **D)** CNN filter depth, **E)** Number of units per dense layer, **F)** Number of dense layers (for both sequence and structure parts).

**Supplementary Figure S2:** Optimization results for hyperparameters for the structure part of the bi-modal CNN. **A)** Batch size optimization, **B)** Dropout rate optimization, **C)** Epoch size optimization **D)** CNN filter depth for the first CNN layer, **E)** CNN filter depth for the second CNN layer, **F)** Number of units per dense layer,

**Supplementary Figure S3:** Optimization of the final output layer **A)** For activation functions, **B)** Learning Optimizers.

**Supplementary Figure S4:** Performance benchmarking result and comparison of DeepPlnc. **A)** The claimed performance by the respective tools, **B)** DeepPlnc’s performance against the same tool when the tool’s datasets were used for training and testing. DeepPlnc outperformed all the tools. C) Initially, on its own dataset PreLnc claimed higher accuracy than DeepPlnc. However, when both the tools were compared for their performance on the remaining four tools’ datasets one-to-one, it emerged that DeepPlnc consistently outperformed PreLnc not only on Datasets A and B, but also on all the four tool’s datasets too.

