## Supplementary Figure S1 for "DeepPlnc: Bi-modal Deep Learning for Highly Accurate Plant lncRNA Discovery"

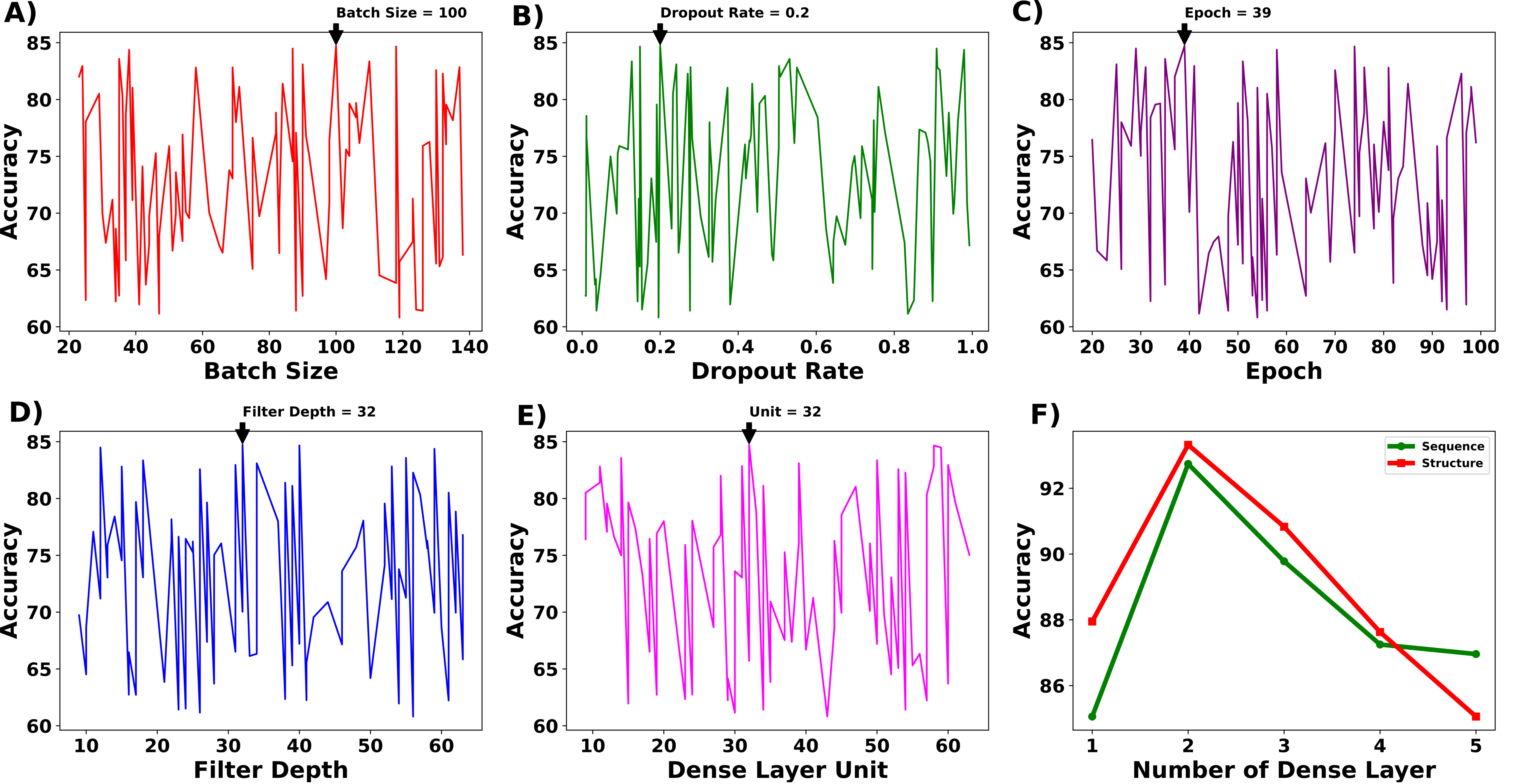


**Supplementary Figure S1:** Optimization results for hyperparameters for sequence part of the bi-modal CNN. **A)** Batch size optimization, **B)** Dropout rate optimization, **C)** Epoch size optimization, **D)** CNN filter depth, **E)** Number of units per dense layer, **F)** Number of dense layers (for both sequence and structure parts).
