## Supplementary Figure S2 for "DeepPlnc: Bi-modal Deep Learning for Highly Accurate Plant lncRNA Discovery"

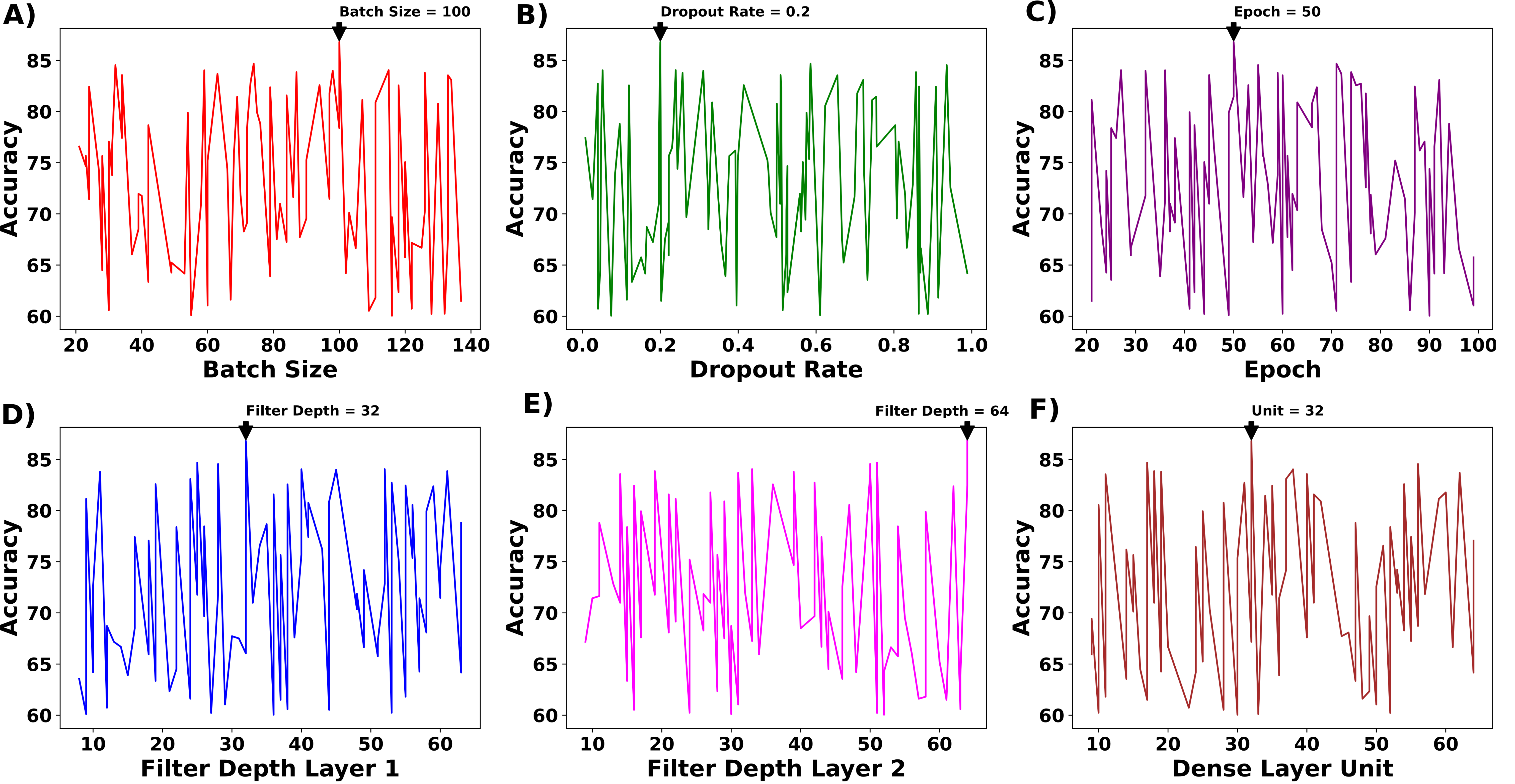


**Supplementary Figure S2:** Optimization results for hyperparameters for the structure part of the bi-modal CNN. **A)** Batch size optimization, **B)** Dropout rate optimization, **C)** Epoch size optimization, **D)** CNN filter depth for the first CNN layer, **E)** CNN filter depth for the second CNN layer, **F)** Number of units per dense layer.
