## Supplementary Figure S4 for "DeepPlnc: Bi-modal Deep Learning for Highly Accurate Plant lncRNA Discovery"

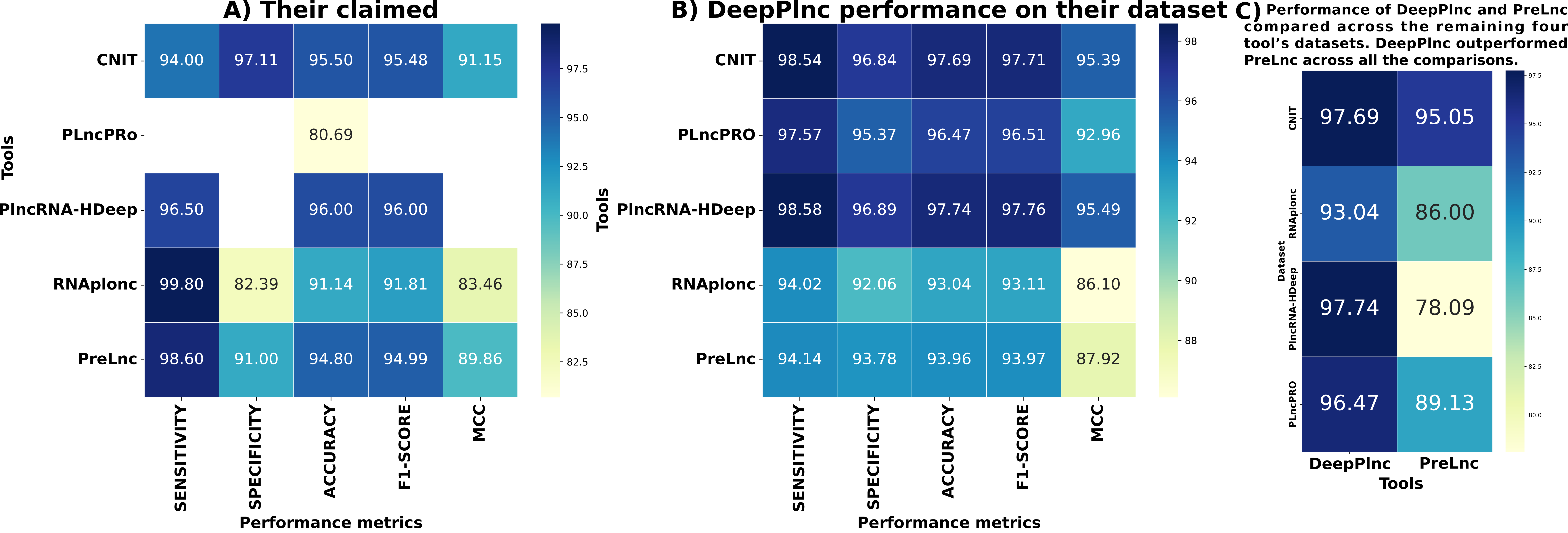


**Supplementary Figure S4:** Performance benchmarking result and comparison of DeepPlnc. **A)** The claimed performance by the respective tools, **B)** DeepPlnc's performance against the same tool when the tool's datasets were used for training and testing. DeepPlnc outperformed all the tools. **C)** Initially, on its own dataset PreLnc claimed higher accuracy than DeepPlnc. However, when both the tools were compared for their performance on the remaining four tools' datasets one-to-one, it emerged that DeepPlnc consistently outperformed PreLnc not only on Datasets A and B, but also on all the four tool's datasets too.
