## Supplementary Figure S3 for "DeepPlnc: Bi-modal Deep Learning for Highly Accurate Plant lncRNA Discovery"

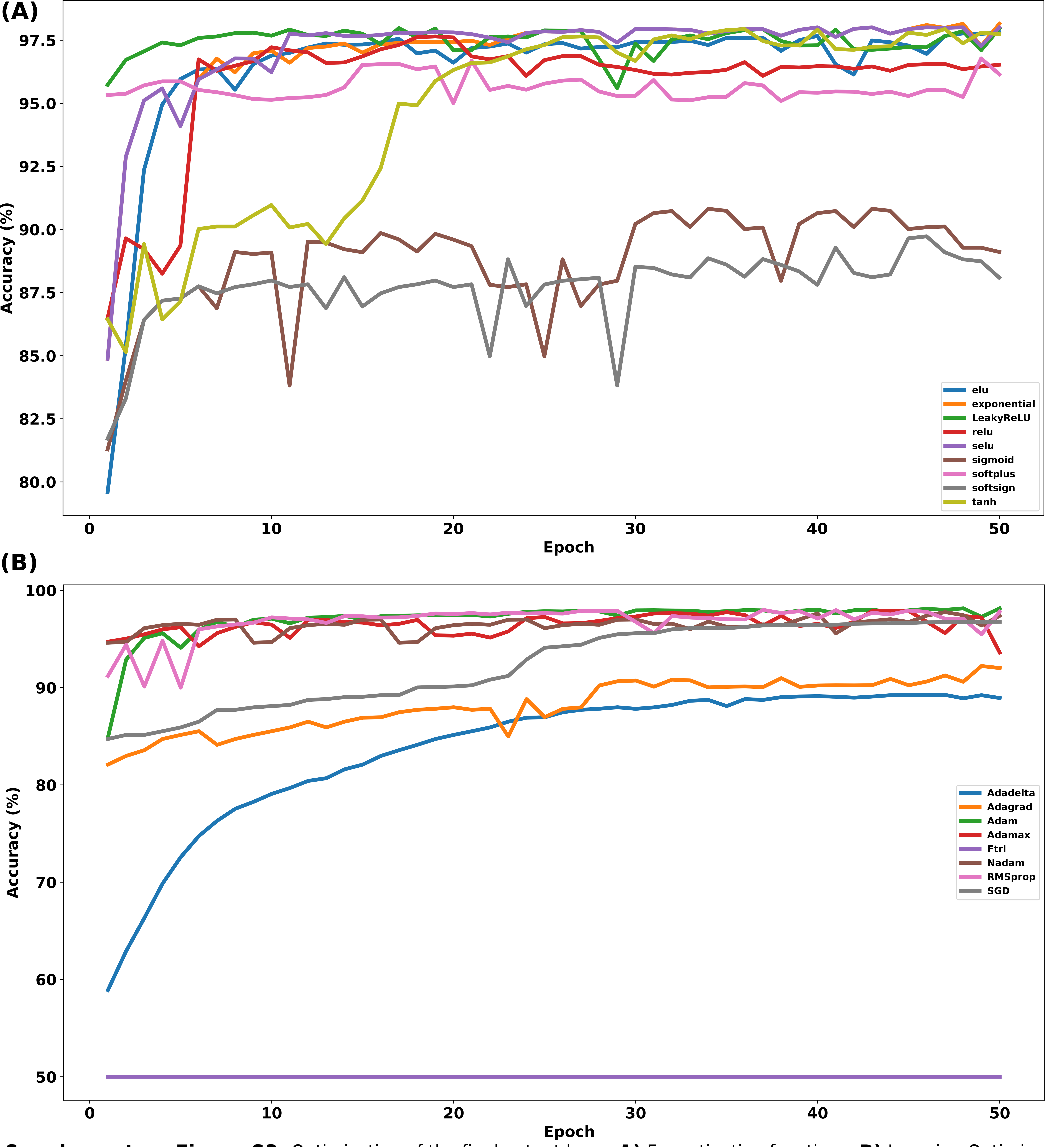


**Supplementary Figure S3:** Optimization of the final output layer. **A)** For activation functions, **B)** Learning Optimizers.
